# Genome-wide analysis of Heavy metal ATPase (P1B-type ATPase) gene family in Mung bean [*Vigna radiata* (L.) R. Wilczek var. *radiata*] and their expression analysis under heavy metal (Zn, Cd and Cu) stress

**DOI:** 10.64898/2026.03.25.713876

**Authors:** Jogeswar Panigrahi, Deeptimayee Panigrahy, Banita Rath, Kapil Gupta

## Abstract

Heavy metal ATPases (HMAs) are important group of transmembrane proteins involved in homeostasis of metal ions in plant systems. In this study, a comprehensive analysis of genome assembly (VC1973A v7.1) resulted in the identification of nine HMA genes (*VrHMA*) and their corresponding proteins in Mungbean, an agronomically important legume crop known for its nutritional values. *VrHMA* proteins were also characterized based on their biomolecular features, conserved domains and motifs arrangement, transmembrane helices, pore-line helices, subcellular location and occurrence of signal peptides. Based on sequence homology, nine *VrHMAs* were clustered into two major substrate-specific groups: *VrHMA1*, *VrHMA5* and *VrHMA7* were categorized under the Zn/Co/Cd/Pb ATPase group, whereas the remaining six *VrHMAs* belong to the Cu/Ag subgroup. Gene structure analysis and promoter scanning revealed the structural divergence and presence of various stress-responsive *cis*-acting elements, respectively. The expression analysis of *VrHMA* genes in root and leaf tissues, in response to heavy metal (Zn, Cd and Cu) stress, indicates their role in the uptake, transport and sequestration of metal ions. Interestingly, *VrHMA5* showed incremental upregulation in roots in response to all three heavy metal stresses, whereas its expression was only upregulated in the leaf tissues under Zn stress, which indicates its role in vascular transport in *V. radiata*. In addition, this study provides valuable insights into the functional roles of *VrHMA* genes and will lay a foundation for future genetic improvement in mung bean aimed at enhanced heavy metal stress tolerance and micronutrient homeostasis.

## Introduction

Mung bean [*Vigna radiata* (L.) Wilczek R. var*. radiata*, Family- Fabaceae] is a prominent grain legume crop in Southeast Asian countries, including India [45]. Mung bean possess a diploid genome with 22 somatic chromosomes, and its genome size was estimated to be 579LMbp [30]. The dehulled seed grains of mung bean have been served along with cereals and is a critical source of proteins and essential minerals in the vegetarian diets across the world, and these grains are rich sources of protein (20–30%), dietary fibres, vitamins, minerals, and large quantities of bioactive components, such as peptides, polyphenols, and polysaccharides [23]. In addition, the plant parts exhibit several bio-medicinal properties, such as anti-inflammatory, antibacterial, and antioxidant etc. [45].

Now-a-days, half of the global population prefers vegetarian diets and several of them suffer from micronutrient deficiency as well as low immunity, retarded cognitive progression, aberrant brain development and child growth etc. [20]. Thus, the edible parts of several food grains need to be biofortified with balanced micronutrients, and it is quite desirable to fight against the malnourishment of micronutrients [83]. The content of these micronutrients in the seed grains and edible plant parts varies from genotype to genotype depending upon the plant type, soil composition, agroclimatic conditions, and the adoption of different mechanisms to maintain the balance of various macromolecules and micronutrients. Among the micronutrients, zinc (Zn), copper (Cu), manganese (Mn) and iron (Fe) are essential for plants, whereas lead (Pb) and cadmium (Cd) are non-essential ones. Both groups of metallic micronutrients are toxic to plant systems vis-à-vis their consumers in the food chain beyond certain limits, whereas their deficit also affects the development of plants, crop yield and the nutritionally balanced diet of consumers. These essential micronutrients, such as Cu, Mn and Zn, are present in various proteins and are also associated with several metabolic processes pertaining to growth and differentiation, plant immunity, cellular processes, signal transduction etc. [83]. Hence, plant systems have evolved several mechanisms to regulate micronutrient uptake by roots, xylem loading and unloading in the roots, long-distance transport through vascular tissues, phloem loading and unloading in the edible parts, and vacuolar sequestration [53, 83]. The members of several gene families, including heavy metal ATPase (HMA), are implicated in these processes.

The members of HMA gene family, also known as P1B-type ATPase under the P1 ATPase superfamily, is one of the group of critical heavy metal ion transporters and they facilitate the root uptake of heavy metal ions from the soil with hydrolysis of ATP, maintain homeostasis in association with other transporters and also efflux the metallic cations from the cytoplasm into vacuoles or to the apoplast [25, 31]. A typical HMA gene comprises three cytoplasmic domains, *viz.* the actuator (A), phosphorylation (P), and nucleotide-binding (N) domains, along with the membranous (M) domain featuring six core transmembrane helices. The P-domain includes a conserved DKTGT motif, where an aspartic acid undergoes ATP-dependent phosphorylation and dephosphorylation during the transport cycle. The N-domain, functioning as an intrinsic protein kinase, binds ATP to facilitate phosphorylation in the P-domain. The A-domain is an inbuilt protein phosphatase, and it possesses the TGE motif responsible for the dephosphorylation event in the P-domain [18]. The transmembrane helix-4 (TM4) of the M-domain contains a proline-rich canonical binding site (CBS), where the ligands (such as Zn^2+^) bind during transportation. Moreover, HMAs possess metal-binding domains (MBDs), typically located in the N-terminus and often present in the C-terminus in several cases [64]. Additionally, the HP locus, PXXK motif, GDGxNDxP motif and a CPx/ SPC motif, essential for heavy metal ion transport, are also present in the members of HMAs. To date, the members HMA gene family have been broadly classified into two categories based on their metal ion specificity, such as Cu/Ag-ATPases and Zn/Co/Cd/Pb-ATPases [15, 75].

The first report on cloning of plant HMA gene (AtHMA6/PAA1) was described in *Arabidopsis thaliana* [65] and since then, the HMA gene families have been characterized at genome level in several plant species such as *A. thaliana, Triticum aestivum, Arachis hypogea, Glycine max, Solanum esculentum, Medicago truncatula*, *Areca catechu, Hordeum vulgare* and *Oryza sativa* due to accumulation of genomic data across time, and were quite assorted in terms of their subcellular localization, substrate specificity and other cellular functions [4, 10, 15, 21, 31, 39, 44, 49, 66, 82]. In model plants, *A. thaliana* (dicot) and *O. sativa* (Monocot), the HMA gene family contains eight AtHMAs and nine OsHMAs, respectively. These members of HMA gene family are classified into distinct subgroups: AtHMA1-4 and OsHMA1-3 were placed in the Zn/Co/Cd/Pb subgroup, whereas AtHMA5–8 and OsHMA4–9 belong to the Cu/Ag subgroup [66].

The decoding of whole genome sequences enables functional genomics studies in several plant systems, including the genome wide analysis of various gene families and their characterization. The decoded draft genome of *V. radiata*, its enrichment and the pan-genome data provide a platform for genome-wide analysis of the gene families as well as other genomic exploration in *Vigna* spp. [19, 30, 42]. Several gene families (OSCA, LEA gene, B-box, HSP70) have already been characterized in *V. radiata* [29, 41, 79, 80]. However, the genomic identification of the members of the HMA gene family in *V. radiata* and their characterization have not been reported till date.

Keeping this into account, we report here the genomic identification of HMA gene family in *V. radiata* (*VrHMAs*), their biochemical characterization and domain/ motifs analysis, elucidation of phylogeny, and differential expression analysis of *VrHMA* genes under heavy metal (Zn, Cd and Cu) stress.

## Materials and Methods

### Genomic identification of V*rHMA* genes and their characterization

The whole genome assembly, gene sequence, CDS and protein sequence of mung bean available in Legume Information System (LIS; *V. radiata* VC1973A v7.1) were downloaded and used in the present study [19]. A total of 30,958 predicted genes and gene products, as well as their chromosomal locations on the genome, were loaded into the local MySQL database created for ease in analysis. The members of the HMA gene family in *V. radiata* were identified by three approaches. First, the HMM profile of the conserved domain of HMA (i.e. E1–E2 ATPase: PF00122; Hydrolase: PF00702; HMA: PF00403) from the Pfam database (hosted by Interpro) was submitted as a query using the HMMER 3.3 programme, and the putative *HMA* proteins and their corresponding genes in *V. radiata* were retrieved with E- value ≤ 10^-3^ [48]. Secondly, the protein database of *V. radiata* was searched through the VignaMine platform of LIS, for the coexistence of three conserved domains (PF0122, PF00702 and PF00403) of HMA proteins. Thirdly, the HMA protein sequences of *A. thaliana* (*AtHMA-1* to *-8*), retrieved from the TAIR database, were compared with proteins of VC1973Av7.1 for the recovery of putative HMAs of *V. radiata*. The findings from these analyses were compared, and the probable *VrHMA* protein sequences were examined using the CD-HIT website and SMART Web server to screen for the coexistence of both E1-E2 ATPase and Hydrolase domains [26, 37]. The gene annotation files for HMA proteins were obtained using the batch sequence search function in the conserved domain database and were used to draw the domain map using TBtools v2.097 [9, 26]. Finally, the nine *VrHMA* genes were mapped to the mung bean genome and were named *VrHMA1 to VrHMA9* in accordance with their location on mung bean chromosomes (*Vradi01* to *Vradi11*) in ascending order [19].

Various biochemical properties of *VrHMA*s, including polypeptide length (aa), molecular weight (MW), isoelectric point (pI), hydropathicity average (GRAVY), aliphatic index and instability index, were determined using the PROTOPARAM tool on Expasy server [17]. The arrangement and occurrence of transmembrane helices (TMHs), pore-lining helices (PLHs), and signal peptides were determined using the MEMPACK tool of the PSIPRED protein analysis suite [52], and the occurrence of TMHs and signal peptides was validated through TMHMM and SignalP 6.0 servers, respectively [33, 54]. The post-translational modification (PTM) signature sequences present in each *VrHMA* protein were also uncovered using ScanProsite [11]. The ProtComp v9.0 programme was used to determine the putative subcellular location of each *VrHMA* proteins [61].

### Analysis of conserved motifs in *VrHMA* proteins

The occurrence of conserved motifs, such as DKTGT, HP CPx/SPC, TGE, GDGxNDxP, and GMxCxxC, in the *VrHMA* proteins was analyzed by using the MAST module of MEME 5.5.7 suite [3] with parameters *viz.* zero or one occurrence per sequence, maximum motif number 13 and motif length 2 to 50. These motifs were annotated using the InterProScan [56], and visualized graphically using Ceqlogo module of MEME 5.5.7 suite.

### Multiple sequence alignment (MSA) and Phylogenetic analysis of *VrHMA* proteins

All nine *VrHMA*s along with eight *AtHMA* proteins were aligned using the multiple sequence comparison by log-expectation (MUSCLE) algorithm in MEGA-X software with default parameters to eliminate redundant *VrHMA* sequences, and also to align the transmembrane domains, showing their position outside or inside of the cell membrane, and other conserved domains/ motifs [48]. The alignment result of MSA was refined and visualized using the GeneDocv2.7.0 tool [50]. Based on their sequence homology and conserved motifs, the intraspecific phylogenetic relationships among these nine *VrHMA* proteins were also delineated.

Subsequently, the non-redundant HMA proteins of four model plant species, viz. *A. thaliana*, *M. truncatula*, *G. max*, and *O. sativa*, were obtained [10, 15, 44, 66] and were aligned along with nine *VrHMA*s as explained above. The interspecific phylogenetic tree was also constructed on the basis of their sequence homology and arrangement of conserved domians and motifs following the neighbor-joining (NJ) method with the default parameters except p-distance model and 1000 bootstrap replications [34].

### Chromosomal distribution of *VrHMAs*, Synteny analysis and KA/Ks calculation

The genomic position of all nine *VrHMA*s on *V. radiata* genome assembly (VC1973A v7.1) has been retrieved, and were depicted in the circos genome map using TBtools v2.097. One step McScanX was set with default parameters to identify the syntenic relationships among *V. radiata*, *A. thaliana*, *M. truncatula* and *G. max*, and their outcomes were graphically represented using multiple synteny plots generated with TBtools v2.097. The substitution rate of non-synonymous (Ka) as well as synonymous (Ks) codons and the substitution ratio (Ka/Ks) were obtained through the Nei-Gojobori model using Mega-X software. The approximate divergence time of each paralogous *VrHMA* gene pair was determined using the formula T (Million years ago; MYA) = (Ks/2λ) × 10^-6^, where ‘λ’ represents the average synonymous substitution rate and is equal to ∼ 6.1 × 10^-9^ [30].

### *VrHMA* gene structure and its promoter analysis

The nine *VrHMA* gene sequences and their corresponding coding sequence (CDS) were submitted to the gene structure display server, GSDS 2.0, along with the phylogenomic dendrogram of the *VrHMA*s to determine the genomic organization of exons, introns and UTRs [24]. Based on the chromosomal location of *VrHMA* genes, the probable promoter sequence (upstream sequences from -2000 to +1 bp) of each *VrHMA* genes was obtained from the genome assembly VC1973A v7.1, and was investigated for the cis-acting elements (CAEs) using the Plant CARE database [36]. Then, the position of some stress-responsive CAEs was graphically represented.

### Expression analysis of *VrHMA* genes *in silico*

The in-silico expression values for seven out of nine *VrHMA* genes were analysed by harnessing the transcript abundance data [in log_2_ transformed transcript per million mapped reads (TPM)], from PlantExp Server (https://biotec.njau.edu.cn/plantExp/), and their expression pattern in different plant parts *viz*. root, leaf, flower, seed and pods were visualized as the heat map.

### Effect of heavy metal stress on expression of *VrHMA* genes by qRT PCR

Seeds of a Mung bean (*V. radiata* cv. Kanica) genotype were surface sterilized with 0.1% mercury chloride and then sown in vermiculite and acid-washed sand mix (2:1) for germination. After three days of germination, three seedlings were transplanted in each plastic pots containing acid-washed sand, and each pot was fed with Murashige and Skoog (MS) salts solution (pH 5.8) as a source of nutrients on every alternate day. The two-week-old seedlings were exposed to heavy metal stress for 72 hours by adding an additional 30µM CdCl_2_, 50 µM CuSO_4_ or 500 µM ZnSO_4_ in MS solution, respectively [44]. A set of plants was left untreated and was considered as control. Three replicates for control and each heavy metal treatment (24, 48 and 72 hours) were tested. Plants were grown in a transgenic house under a 14-hour photoperiod, including sunlight irradiance, 27 ± 2°C temperature, and 65-75% relative humidity. Immediately after 24, 48 and 72 hours of treatment, the young roots and apical leaves were sampled under liquid nitrogen and stored at -80°C for RNA isolation and subsequent gene expression analysis. Total RNA was isolated and purified from 50 mg of tissues using the plant RNA isolation kit (gene2me, India) following the manufacturer’s instructions. The assessment of RNA quality and yield was made using Qubit 4 Fluorometer (Invitrogen, USA), and was re-affirmed by agarose gel electrophoresis. One microgram of purified RNA was used for the synthesis of cDNA, expending 2 µl of Verso enzyme mix (200 Ul^-1^), oligo(dT) primer (1.0µg) and dNTP mix (5 mM each) following the manufacturer’s instructions (Verso cDNA synthesis kit; Thermo Scientific, USA). The quality and integrity of each cDNA sample were assessed by amplification of the tubulin (*VrTubulin*, Gene ID: LOC106775745) gene using G9 Taq DNA polymerase master mix (GCC Biotech Pvt. Ltd., India) and primers (F: 5’GCATTGGTACACTGGAGAAG3’ and R: 5’TCCTCATACTCGTCGTCATC3’ [68]. Dilution of cDNA samples (1:10) was carried out using nuclease-free water, and 2 µl of the diluted cDNA sample was used for quantitative real-time PCR (qRTPCR) using DyNAmo Flash SYBR green qPCR kit (Thermo Scientific, USA) and the primers depicted in **Supplementary Table 1.** The qRT-PCR assay of 10µl, containing 2µl of cDNA, 5µl of DyNAmo Flash SYBR green mix, 0.5µl of both forward and reverse primer and 2µl of nuclease-free water, was made in an LC-480 type 96-well plate (Tarson, India) using a Real-Time PCR system (CFX Connect, Biorad, USA) following the PCR conditions as described by[48]. Normalization of the expression of *VrHMA* genes, in both control and treated samples, was carried out using *VrTubulin* as an endogenous control. Each analysis was performed with three biological and three technical replicates, and the specificity of amplicons was affirmed by melting curve analysis. The relative expression of *VrHMA* genes was calculated employing 2^-ΔΔCT^ method [43]. The significant difference in expression values was evaluated with one-way analysis of variance (ANOVA) at p=0.05 level of significance using SPSS V 16.0.0 software.

## Results

### *In-silico* identification of HMA gene family in *V. radiata* and their characterization

A genomic HMM search using the three HMA-specific domains, E1–E2 ATPase (PF00122), hydrolase (PF00702) and HMA (PF00403), as query against the 30958 genes of Mung bean genome (VC1973A v7.1) identified only eight non-redundant proteins containing all three domains with e-value 10^-3^ or less, whereas 38 proteins containing both E1–E2 ATPase and hydrolase domains. Similarly, the search through Vigna Intermines resulted in five non-redundant proteins containing all three domains, whereas 38 proteins had both E1–E2 ATPase and hydrolase domains. Further, the homology search of HMA proteins from *A. thaliana* (*AtHMA1-8*) resulted in nine putative HMAs with e-value 10^-3^ or less. After combining the findings of three approaches, a total of 38 probable proteins with both E1–E2 ATPase and hydrolase domains were further characterized for the occurrence of conserved domains/motifs, and only nine out of 38 possessed conserved motifs unique to HMA members of model dicots, and were considered as members of the HMA gene family of *V. radiata* (*VrHMAs*). The gene sequence of nine *VrHMAs* were present on five different chromosomes (Vradi-02, -03, -08, -09 and -11) of the *V. radiata* genome (**Table 1**) and were named *VrHMA1* to *VrHMA9* considering their location on different chromosomes in ascending order. The size of *VrHMA* proteins varied from 781 to 1010 aa, and their corresponding molecular weight also ranged between 84.7 kDa (*VrHMA4*) to 109.1 kDa (*VrHMA6*), with an average value of 96.4 kDa **(Table 1)**. All nine *VrHMA*s commence with an initiation codon and terminate with a stop codon as expected, and their theoretical pI values were estimated to be within the range of 5.57 (*VrHMA2*) to 7.53 (*VrHMA1*) with an average of 6.26. Based on amino acid composition, all the *VrHMA* proteins were hydrophobic in nature with positive GRAVY value (0.056–0.299). The aliphatic index of VrHMA proteins was in the range of 99.95 to 105.01, with a higher amount of aliphatic amino acids, and are likely to be stable across a wide range of temperatures. The instability index of all nine *VrHMAs* was also less than 40 (33.73 to 38.57), which indicates their high stability *in vitro*. The subcellular locations of these nine *VrHMAs* were predicted to be either in the plasma membrane or the chloroplast **(Table 2)**.

**Table 1:**
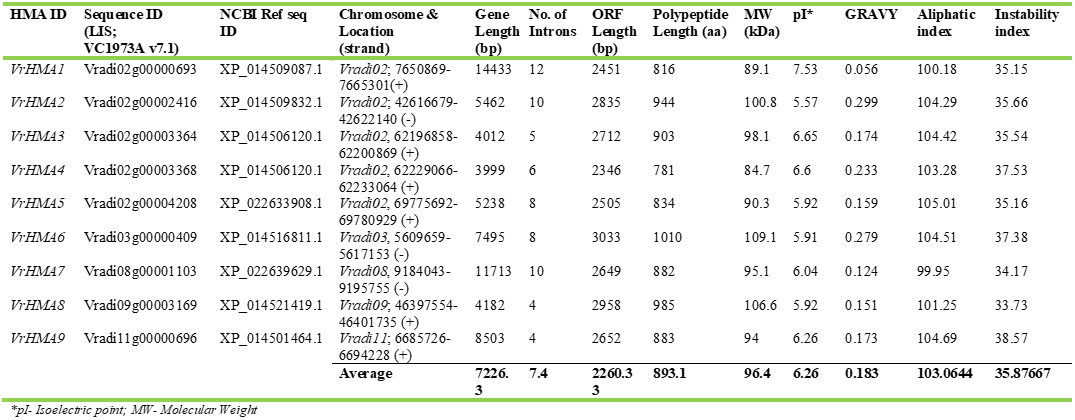
Biomolecular properties of nine *VrHMA* genes identified in Mung bean *(VrHMA)* and their proteins.

**Table 2:**
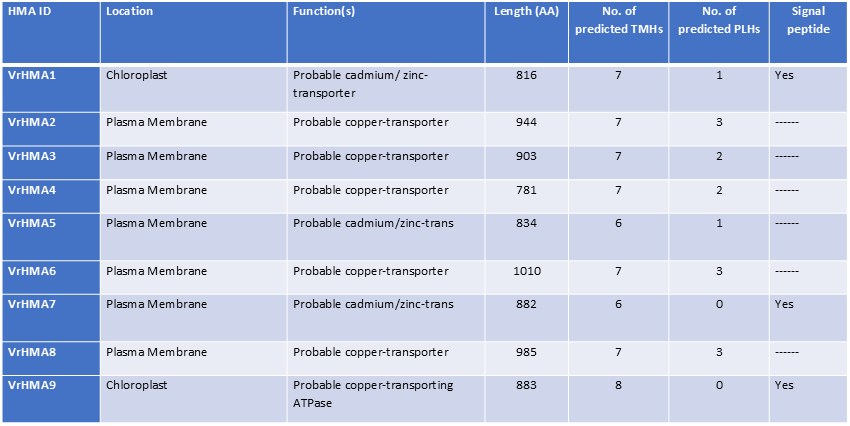
Subcellular localization and functions of nine *VrHMA* proteins along with their transmembrane helices (TMHs), pore lining helices (PLHs) and signal peptide sequences.

### Intra- and Inter-specific phylogenetics analysis of VrHMAs

The MSA of nine *VrHMA* protein sequences were used to create an unrooted intraspecific phylogenetic tree, which revealed that these *VrHMA*s were grouped into two major clusters depending upon their substrate specificities as predicted in terms of their functions **(Table 2)**. Three *VrHMAs* (*VrHMA1*, *VrHMA5* and *VrHMA7*) belong to the Zn/Co/Cd/Pb ATPase subgroup, whereas the remaining six *VrHMA*s belong to the Cu/Ag ATPase subgroup **(Fig. 1).**

The interspecific phylogenetic relationship of 55 HMAs from *A. thaliana*, *O. sativa*, *G. max, M. truncatula* and *V. radiata*, as delineated by MEGA-X, revealed the clustering of these *VrHMA* proteins into six clades (I–VI), and were further sorted into two major groups representing divalent Zn/Co/Cd/Pb-ATPases (clade I and II) and monovalent Cu/Ag-ATPases (clade III-VI) based on their specificity and functions as denoted above **(Fig. 1).** Although the numerical distribution of *VrHMAs* into distinct groups was not equal, the clustering of *VrHMA* proteins into the major groups, based on their substrate specificity, along with the corresponding HMA proteins of *A. thaliana* and *M. truncatula*, was quite unambiguous. For example, the clade-I and II contains three members of *VrHMA* gene family (*VrHMA1*, *-5* and *-7*) along with *AtHMA1-4* (4), *MtHMA2* and -6 (2), *GmHMA-5, -13, -14, -16, -18* and *-19* (6), and *OsHMA1-3* (3) which are determined to be Zn/Co/Cd/Pb-ATPases type **(Fig. 1)**. The remaining six members of *VrHMA* gene family were unequally distributed among the four clades along with other members of Cu/Ag ATPase group.

### Genomic location, synteny and duplication analysis of VrHMAs

The nine *VrHMAs* were distributed among five chromosomes (*Vradi-02, -03, -08, 09* and *-11*) of Mung bean genome, and five of them (*VrHMA1*-*5*) were located on chromosome *Vradi02*, whereas rest four *VrHMAs* were distributed among four chromosomes, *Vradi-03*, -*08*, *-09* and *-11* **(Fig. 2)**. The intraspecific synteny analysis delineated the possible gene duplication events among the members of the HMA gene family during the evolution of the *V. radiata* genome, and 10 pairs of syntenic *VrHMA* genes were noticed in *V. radiata* **(Fig. 2)**. Among them one gene pair i.e. *VrHMA3/VrHMA4* underwent tandem duplication. Rest nine gene pairs *VrHMA2/VrHMA3*, *VrHMA2/VrHMA6*, *VrHMA2/VrHMA8*, *VrHMA3/VrHMA6*, *VrHMA3/VrHMA8*, *VrHMA4/VrHMA6*, *VrHMA4/VrHMA8*, *VrHMA6/VrHMA8*, *VrHMA5/VrHMA7* underwent segmental duplications. To understand the divergence process of homologous *VrHMA* genes, non-synonymous and synonymous substitutions, and their ratio were calculated among *VrHMA genes* (**Supplementary Table 2**). The Ka/Ks ratio of duplicated gene pairs were ranged from 0.092 to 0.673, which indicates the role of purifying or negative selection during the evolution and divergence of *VrHMA* genes. Interspecific synteny analyses of *VrHMA* genes with the members of HMA gene family of model dicot (*A. thaliana*; Brassicaceae) as well as model legumes (G*. max* and M. *truncatula*; Fabaceae) showed sharing of several gene and genomic segments **(Supplementary Table 3)**. For example six homologous gene pairs (*PAC:19650287/ VrHMA8*; *PAC:19641614/ VrHMA7*; *PAC:19643720/ VrHMA7*; *PAC:19650287/ VrHMA3*; *PAC:19648322/VrHMA1*; *PAC:19665627/ VrHMA9*) was identified between *V. radiata* and *A. thaliana* **(Fig. 3)**. Similarly, 8 and 19 homologous gene pair between *V. radiata* and *M. truncatula*, and *V. radiata* and *G. max* were identified **(Fig. 3; Supplementary Table 3)**.

### Analysis of conserved domains and motifs in the protein architecture of *VrHMAs*

MSA of all eight *AtHMA* and nine *VrHMA* protein sequences revealed that all *VrHMAs* possessed the TGE, DKTGT, PxxK, HP and CPx/SPC conserved motifs unequivocally. But, the TGE motif showed positional alternation in *VrHMA4* and *VrHMA5*. Similarly, the GDGxNDxP motif was present in all *VrHMAs* except *VrHMA1*, where the GEGxNDxP motif was present in lieu of GDGxNDxP motif. Similarly, the P1B type-specific CPx motif was present in all *VrHMAs* except *VrHMA1*, where CPx is replaced by the SPC motif **(Fig. 4).** The GMxCxxC motif, a unique metal binding motif for the Cu/Ag-ATPase group, was present in all six *VrHMAs* of Cu/Ag ATPase group (*VrHMA-2*, *-3*, *-4*, *-6*, *-8* and *-9*), whereas *VrHMA5* and *VrHMA7*, belong to Zn/Co/Cd/Pb ATPase group, possessed with two cysteine residues of the GxCCxxE motif. But, the *VrHMA1* had N-terminal histidine-rich (His-rich) motif in lieu of GxCCxxE motif **(Fig. 4).**

CDD analysis reaffirmed the presence of three characteristic domains, viz. hydrolase, heavy metal associated (HMA) and E1-E2 ATPase domain in all *VrHMAs* except *VrHMA1*, where only hydrolase and E1-E2 ATPase were present **(Fig. 5a).** The same distribution pattern of domains was also evident through SMART web server **(Supplementary Fig. 1).** Further analysis of motif diversity among nine *VrHMAs* using MEME suite revealed the presence of conserved motifs such as DKTGT, GDGxNDxP, HP, TGE, CPx/ SPC and GMTCxxC motifs **(Fig. 5b)**. Further, the functional annotation of 13 motifs through Interproscan revealed that most of them belong to P-type cation transporting ATPases **(Supplementary Table 4).** The sequence logo generated through Ceqlogo also revealed conserved motifs in *VrHMAs* as expected **(Supplementary Fig. 2)**. The comparative analysis also revealed that all *VrHMA*s except *VrHMA*1 contain motifs 1, 2, 5, 7, 8, 9 and 12 which represent the important motifs of P1B type ATPase proteins. In *VrHMA*1, the motif 12 representing metal binding motif (GMTCxxC) was absent, and also variations in the location of other motifs across different *VrHMA* genes were also noticed.

The MEMPACK of PSIPRED, TMHMM 2.0 and SignalP 4.1 server outlined the number of TMHs, PLHs and signal peptides present in the *VrHMA* proteins **(Table 2**; **Fig. 6)**. Like other species, *VrHMA* genes possessed with six to eight TMHs, and were embedded with varying numbers (one to three) of PLHs to facilitate passive and active transport of ions across the membranes. Moreover, three *VrHMAs* (*VrHMA1*, *VrHMA7* and *VrHMA9*) possessed signal peptide sequence, and two of them (*VrHMA1* and *VrHMA9*) were predicted to be in chloroplast and the remaining seven *VrHMA* proteins were located in the plasma membrane **(Table 2).**

The *VrHMA* proteins were predicted to have several post-translational modifications (PTM) peptide sequences, such as N-glycosylation site, protein kinase-C (PKC) phosphorylation site, N-myristylation, P-type ATPases phosphorylation site and heavy-metal-associated domain related sites etc. **(Table 3)**. Among the observed PTMs, the N-myristylation peptide sequence was predominantly present in all *VrHMA*s (10 in *VrHMA1* to 23 in *VrHMA2*) and was followed by PKC phosphorylation, casein kinase-II phosphorylation and N- glycosylation sites, respectively. All *VrHMA* proteins, except *VrHMA1*, possessed with atleast a PTM site related to the heavy metal-associated domain, and in lieu of it the *VrHMA1* possessed with a histidine-rich region. The *VrHMA7* also had the cysteine-rich region as PTM sequence **(Table 3).**

**Table 3:**
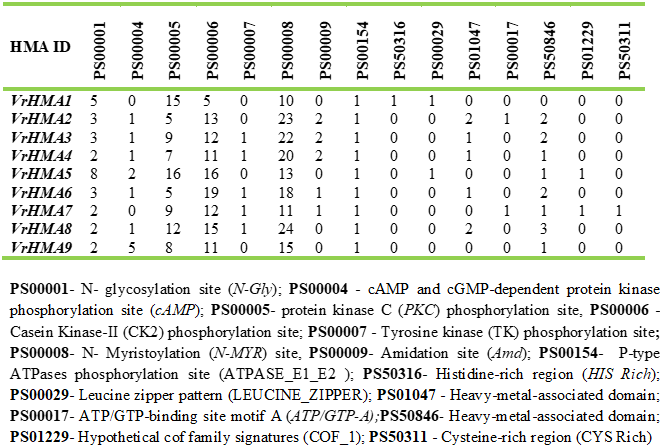
Post-translational modification sites predicted in nine *VrHMA* proteins of Mung bean.

### Analysis of gene structure and promoter sequence of *VrHMAs*

Gene structure analysis of nine *VrHMAs* on GSDS 2.0 revealed variation in terms exon-intron arrangement, their number and length **(Supplementary Fig. 3).** The length of nine *VrHMA* genes was found to be assorted between 3999 bp (*VrHMA4*) and 14433 bp (*VrHMA1*), and the length of CDS was ranged between 2346 bp (*VrHMA4*) and 3033 bp (*VrHMA6*). The number of introns also varied from four (*VrHMA8* and *-9*) to 12 introns (*VrHMA1*). The number of introns also varied considerably within both the Cu/Ag ATPase (4-10) and Zn/Co/Cd/Pb ATPase (8-12) groups of *VrHMAs*. This variation in structural arrangement of *VrHMA* genes was well imitated in the MSA among the members *VrHMA* proteins **(Fig. 4)**, whose length varied between 781 and 1010 aa with similarity ranging from 17.76% (*VrHMA5* and *VrHMA9*) to 61.37% (*VrHMA3* and *VrHMA8*) (**Supplementary Table 5).**

Analysis of the promoter sequences of nine *VrHMA* genes using PlantCARE database revealed the presence of various stress responsive *cis*-acting elements (CAEs), such as ABA-responsive elements (ABREs and AREs), MeJA-responsive motifs (CGTCA or TGACG), W-Box, WUN motif, ethylene responsive elements (EREs), TC-rich element, light regulatory element (LREs), low temperature responsive elements (LTRE), Gibberellic acid responsive elements (GARE), light-responsive G-boxes and W-boxes **(Supplementary Table 6; Fig. 7)**. The promoter of *VrHMA3* contains maximum number (204) of CAEs, whereas *VrHMA1* has 135 CAEs. MeJA-responsive CGTCA motifs or TGACG motifs present in *VrHMA1*, *VrHMA2*, *VrHMA7* and *VrHMA8*. The *VrHMA-2*, *-3*, *-4*, *-5*, *-8* promoters contain TC-rich repetitive elements involved in defense and stress responses. The ABRE element present in all *VrHMAs* except *VrHMA4*. The *VrHMA1*, *VrHMA6*, *VrHMA7* and *VrHMA8* contain TCA elements which are related to salicylic acid response. The *VrHMA1*, *VrHMA3* and *VrHMA9* promoters contain low-temperature responsive (LTR) CAEs. The GARE motif, which shows Gibberellic acid specific responses, was present only in *VrHMA2* and *VrHMA8*, and the auxin-responsive TGA element was only present in *VrHMA2*, *VrHMA6* and *VrHMA7*.

### Expression analysis of *VrHMA* genes *in silico*

Tissue specific expression analysis of seven out nine *VrHMA* genes (except *VrHMA3 and VrHMA4*), in terms of transcript abundance (TPM value) in five tissues, were varied from tissue to tissue showing quite heterogeneous expression pattern **(Fig. 8)**. Expression profiling revealed that four genes (*VrHMA1, -2, -8*, and *-9*) showed higher expression in root tissue, whereas rest three *VrHMAs* showed low to moderate level expression. In leaves, four *VrHMA* genes (*VrHMA1, -6, -7,* and *-9*) showed higher expression, whereas the remaining three showed moderate to low level expression. In flowers, all seven *VrHMA* genes showed higher expression except *VrHMA5*, whereas in seeds, five *VrHMA* genes (*VrHMA2, -5, -6, -7*, and *-9*) showed meagre to no expression, and the remaining two genes also showed very low-level expression. In pods, three *VrHMA* genes (*VrHMA1, -7*, and *-8*) showed higher expression, whereas three *VrHMAs* (*VrHMA-2, -6*, and *-9*) showed moderate to low level expression, and *VrHMA5* showed meagre expression.

### Expression analysis of *VrHMAs* under heavy metal stress induction

The heavy metal (Zn^2+^, Cd^2+^ and Cu^2+^) induced response of *VrHMA* genes in roots and leaves of a Mung bean genotype (*V. radiata* cv. Kanica) was assessed. In response to Zn^2+^ stress, expression of all *VrHMA* genes were upregulated in roots, either gradually with the increment of time up to 72 hours or after a specific duration (24 hours or 48 hours) of treatment **(Fig. 9a)**. In contrast, eight out of nine *VrHMA* genes (except *VrHMA3*) also showed upregulated expression in leaves **(Fig. 9b)**. In both roots and leaves, two (*VrHMA1* and *VrHMA5*) out of three *VrHMAs* belong to the Zn/Co/Cd/Pb-ATPases subgroup and showed incremental upregulation in this study, whereas *VrHMA7* showed incremental upregulation only in roots. In leaves, *VrHMA6* and *VrHMA7* were upregulated (∼ 10-fold) after 24 hours of treatment, and thereafter gradually downregulated with time **(Fig. 9a, b)**. The expression of *VrHMA8* was upregulated after 24 hours in both roots and leaves, and then declined gradually with time in roots, but showed incremental upregulation in leaves up to 72 hours. *VrHMA4* showed the highest upregulation (∼ 30 fold) in roots after 24 hours of Zn stress, and in leaves *VrHMA5* showed highest upregulation (∼30 fold) after 48 as well as 72 hours of treatment.

In response to Cd^2+^ stress, eight out of nine *VrHMA* genes showed heterogeneous overexpression both in roots (except *VrHMA1*) and leaves (except *VrHMA5*), either gradually with the increment of time or after a specific duration (up to 72 hours) of treatment **(Fig. 9c, d)**. Like Zn^2+^ response, *VrHMA5* and *VrHMA7*, which belong to Zn/Co/Cd/Pb-ATPases subgroup, showed incremental upregulation in roots in response to Cd^2+^ stress, whereas *VrHMA1* was not expressed even after 72 hours of Cd stress treatment **(Fig. 9c)**. The expression of *VrHMA7* was upregulated after 24 hours in both roots and leaves, and then showed incremental upregulation in roots, but downregulated in leaves up to 72 hours of the treatment. In both roots and leaves, *VrHMA2* and *VrHMA4* showed the highest upregulation (15-30 fold) after 24 hours of Cd stress, and in roots, *VrHMA5* showed highest upregulation (∼30 fold) after 72 hours of treatment **(Fig. 9d)**.

In response to Cu^2+^ stress, five out of nine *VrHMA* genes (*VrHMA1, -4, -6, -7* and *-8*) were overexpressed in roots after 24 hours of treatment and then down-regulated gradually with the increment of time **(Fig. 9e)**. In contrast, five out of nine *VrHMA* genes (*VrHMA1, -2, -3, -8* and *-9*) showed upregulated expression in leaves with increment of time upto 72 hours of treatment **(Fig. 9f)**. The *VrHMA5* showed consistent upregulation in roots after both 48 and 72 hours of treatment, whereas it was not expressed in leaves.

## Discussion

Mung bean grows in diverse soil conditions, including heavy metal-contaminated soils and marginal lands across tropical and subtropical countries of the world. Thus, it possesses several mechanisms regulating absorption of micronutrients (Zn, Fe, Cu, Mn etc.) from the soil, and their transport, distribution and accumulation in the edible parts within a limit, and also absorption and sequestration of nonessential micronutrients (Cd, Pb etc.) as well as the essential micronutrients beyond the toxic limit [31]. Several transporter gene families mediate homeostasis of these micronutrients, and among them, the transmembranous HMAs play a critical role in crop plants [75]. The composition of HMA gene family in the plant systems are quite variable, such as eight in *A. thaliana*, nine in *O. sativa* and *M. truncatula* 31 in *B. napus*, 20 in *G. max,* 21 in *H. vulgare,* 27 in *T. aestivum*, *and 48 in S. lycopersicum* [4, 10, 15, 40, 44, 49, 66, 82]. In this study, nine *VrHMA* genes were identified in *V. radiata* with an estimated genome size of nearly 579 Mbp [30]. The number of *VrHMA* genes in this crop was almost *at par* with those of model dicot (*A. thaliana*), as well as model legume (*M. truncatula*), whose genome sizes were estimated to be ∼125Mbp and ∼465Mbp, respectively [69, 70]. The number of the HMA genes in *G. max*, *T. aestivum*, *H vulgare* and *B. napus* were comparatively higher with estimated genome sizes of ∼1.01 Gbp, 14.6 Gbp, 5 Gbp and 976 Mbp, respectively. However, the comparative analysis revealed there is no correlation between the members of the HMA gene family with genome size, and this variation might be attributed to genetic traits and driving forces in the evolution of individual genomes and genetic systems across time [8, 15].

The members of the HMA gene family clustered into varying number of clades/ groups and subgroups in different plant species based on their substrate specificity and sharing of domains and motifs. HMA proteins in *A. thaliana*, *M. truncatula* and *G. max* were grouped into six clades [10, 15, 44], whereas five in *H. vulgare* and *B. napus* [40, 82], four in *T. aestivum* [4] and 12 in *S. lycopersicum* [49]. In *V. radiata*, nine *VrHMAs* were clustered into six clades based on sequence homology vis-à-vis sharing of domains and motifs *inter se*, and these six clades were further classified into two major groups representing divalent Zn/Co/Cd/Pb-ATPases (clade I and II) and monovalent Cu/Ag-ATPases (clade III-VI) based on their substrate specificity and phylogenetic clustering along with the homologous HMA proteins of *A. thaliana*, *M. truncatula*, *G. max* and *O. sativa*. The clustering of *VrHMA* proteins along with the corresponding orthologous HMA proteins indicates the taxonomic conservation of the structural organization of HMA proteins at the family level as reported for Zn/Co/Cd/Pb P1B ATPase group [84]. Further it has also observed that the structural organization of *VrHMA* proteins, including domains and motifs arrangement, modulated by gene structure and exon-intron arrangement influences phylogenetic topology as well as functional divergence among the members of same gene families, in particular, the transport of specific substrates as reported in several legumes [15, 31, 44].

The HMA proteins with similar architecture of domain and motif arrangement might have similar functions, in particular, the transport of specific substrates as reported in the *MtHMAs* [44]. In the present study, three characteristic domains of HMA proteins, viz. hydrolase, heavy metal associated (HMA) and E1-E2 ATPase domains, were present in all *VrHMAs* except *VrHMA1*, where only conserved hydrolase and E1-E2 ATPase domains were present. A similar observation was also reported in *T. aestivum* [4], and this could be attributed to their multi-functional role in regulating the uptake, vascular movement and release of heavy metal ions through phosphorylation, and provision of energy for membrane transport. Ideally, HMA in plant systems contain eight TMHs [64], and the *VrHMA*s contain six to eight TMHs as expected. These increased or decreased number TMHs in *VrHMA* proteins could be attributed to genomic reorganization during the divergence of HMA gene family [13], and similar variation in the number of TMHs had also been reported in peanut and soybean [15, 39]. One to three PLHs were embedded in the TMHs of *VrHMA* proteins, which might be involved in the perception and modulation of ionic movement across the membranes in response to cellular signaling of heavy metal stress [48].

MSA analysis showed that the *VrHMA* proteins possess six conserved motifs *viz*., TGE, DKTGT, CPX/SPC, GDGxNDxP, HP, and PxxK, unique to P1B-type ATPase proteins. Among them, TGE motif exhibits positional variation by occupying at the N-terminal cytoplasmic side of *VrHMA4* and the extracellular loop connecting the TMH III and IV in *VrHMA5*. In previous studies in *A. thaliana*, mutation in the TGE and DKTGT (phosphorylable D-residues) motifs severely compromises the metal ion transport ability in *ran1-1* mutant, and also the activity of AtHMA3/AtHMA4, respectively [22, 75]. The positional alteration of the TGE motif in two *VrHMA*s might have divergent functions in *V. radiata*. Even in *VrHMA1*, GDGxNDxP and CPx motifs were replaced by GEGxNDxP and SPC motifs, respectively, similar to *AtHMA1*, *OSHMA1*, *GmHMA5* and *GmHMA19* [15, 75]. In addition, the GMXCXXC motif, unique to Cu/Ag type ATPase, was present in all six *VrHMAs* of Cu/Ag ATPase group, whereas the members of Zn/Co/Cd/Pb group except *VrHMA1*, contain GXCCXXE motif. In lieu, *VrHMA1* contains an N-terminal His-rich motif similar to *AtHMA1*, and probably the His-rich motif might be involved in the coordination of divalent cation transport activities in this species [60]. Even in *A. thaliana*, the Zn^2+^ commonly binds to the Cys and His residues, and Cys-His rich motifs present in *bZIP19* and *bZIP23* recruit Zn^2+^ in coordination with cellular Zn status [28]. In *AtHMA2, AtHMA3* and *AtHMA4* have a variant GxCCxxE motif in N-terminal metal binding domain, and the two *Cys* residues present in this motif bind divalent cations (Zn^2+^ and Cd^2+^) with elevated affinity [75]. Either the mutation in Cys residues or truncation of N-terminal metal binding domain causes drastic reduction in ATPase activity and also abolished its ability to complement Zn and Cd hypersensitivity [74]. It had also been reported that a novel zinc binding protein interacts with *Cys* residues (CxxC) through thiol groups and regulate divalent metal ion homeostasis [16]. In addition, several new motifs (3, 4, 13), having P-type ATPase activities as predicted function, were also consistently present in the N- and C- terminal region of *VrHMAs* in the similar arrangement, and probably these motifs are also assisting metal ion homeostasis [31].

The PROSITE analysis revealed numerous phosphorylation sites in *VrHMA*s, which indicates their role as substrates for kinases such as casein kinase II, PKC, P-type ATPases, consistent with their roles in metal homeostasis under abiotic stress across legumes [62, 71]. Notably, N-myristylation was the most prevalent PTM among *VrHMAs*, which is likely contributing to conformational stability under abiotic stress and functions like signal transduction, intercellular export, and zinc/heavy metal membrane transport, as observed in other plants [48, 72, 81]. Even all *VrHMAs* have atleast one P-Type ATPase phosphorylation site, and this site typically involves a conserved aspartate residue within a DKTGT motif, where the aspartate is transiently phosphorylated during ATP hydrolysis to drive metal ion transport [64]. These features highlight critical roles of *VrHMA* proteins in cellular pathways regulating metal homeostasis. The HMA domain-related PTM sites were found to be present in all *VrHMAs* except *VrHMA1*, where the alternate *His*-rich region was present. Posttranslational modifications in these sites might be helping in the metal binding process [18]. Similar to earlier reports, the cys-rich motif present in *VrHMA7* might have some role in sequestration and detoxification of heavy metal through Metallothionein binding [7].

It has been reported that HMAs have distinct sub-cellular localization in model plants and were linked to their transporter activities [6, 46, 58]. In this study, *VrHMA1* the homolog of *AtHMA1*, was also located in the chloroplast similar to *AtHMA1, AtHMA6, AtHMA8, MtHMA4* and *OsHMA1*, which indicates the association of *VrHMA1* with zinc detoxification through vacuole alike *AtHMA1* as well as Zn and Cu translocation into the chloroplast [5, 32, 44, 59, 66]. Even, the chloroplast located members might have been playing significant role in ion delivery into chloroplast to mediate different enzymatic reactions within it [59]. In this study, seven *VrHMAs* (*VrHMA2-8*) were predicted to be located in the plasma membrane, similar to *AtHMA2, AtHMA4, AtHMA5, MtHMA1-3, MtHMA5-9, OsHMA2, OsHMA5* and *OsHMA9*. [2, 12, 14, 35, 44, 46, 67]. This plasma membrane-localized *VrHMAs* might have a role in the root-to-shoot translocation of the heavy metals through xylem loading as reported in *A. thaliana* and *O. sativa* [2, 12, 67, 76]. Moreover, three *VrHMAs* (*VrHMA1, VrHMA7* and *VrHMA9*) possessed signal peptide sequences, and two of them (*VrHMA1* and *VrHMA9*) were predicted to be in chloroplast, which indicates the contribution of these *VrHMA* proteins in the cellular transport of heavy metal ions [31, 39, 44]. Further transformation and complementation studies will delineate the exact location and function of these *VrHMAs* in subsequent studies.

The number of exons and introns also varied among the *VrHMAs* even within substrate-specific clusters (Cu/Ag- and Zn/Co/Cd/Pb ATPase). These variations in exon count and exon-intron arrangements among the members of *VrHMAs* suggest genomic reorganization and potential functional diversification of this gene family during the evolution and domestication of *V. radiata,* as reported in *G. max* and *A. hypogea* [15, 39, 77]. The CAEs are regulatory sequences and they play a significant role during the growth and development of the plant system upon exposure to heavy metal stresses [57]. Promoter analysis of *VrHMAs* also revealed the prevalence of several heavy metal responsive CAEs like MYB, Myc and STRE in addition to other cellular and molecular CAEs, which indicates the significant contribution of *VrHMA* proteins to the plant’s defense against zinc, cadmium, copper and other heavy metals [63]. Several multi-stress responsive CAEs, such as G-box and ABRE, also present in *VrHMA* promoters which might have also played vital role in plant stress responses [31]. The prevalence of these CAEs indicates the key role *VrHMA* genes under diverse abiotic stresses, potentially enhancing the plant’s overall resilience against stress alike other members of Fabaceae [15, 39, 44].

Homeostasis of heavy metals is well modulated by coordinated expression of group of membrane transporters in the plant systems, and among them, HMAs play a crucial role in root uptake and detoxification of heavy metals, either inhibiting influx into the cytoplasm or effluxion from cytoplasm to vacuoles/ apoplasts, in model plants [27, 75]. In silico analysis of RNAseq data (https://biotec.njau.edu.cn/plantExp server) revealed heterogeneous expression of the *VrHMA* genes except *VrHMA3* and *VrHMA4* in five tissues (root, leaves, flower, pod seed). The qRT PCR analysis showed differential expression pattern of these genes in response to heavy metal (Zn^2+^, Cd^2+^ and Cu^2+^) stress, which was indicative of diverse function of these genes as reported in S. *lycopersicum*, A*. hypogaea* and *A. catechu* [31, 39, 49]. Even though the *VrHMA* genes (1, 5 and 7), grouped under Zn/Co/Cd/Pb-type ATPase, showed varied expression in response to Zn and Cd stress, which can be attributed to dissimilar sensing and signaling mechanisms available in *V. radiata* [55]. The expression of *VrHMA5* was upregulated in root tissues in response to all three heavy metal stresses, whereas it showed higher expression in leaf tissues only in response to Zn^2+^ stress. This suggests the role of *VrHMA5* in transport of at least three heavy metal ions in root tissues, and putatively it plays a significant role in either mitigating the toxicity of zinc by sequestering them into vacuoles or transporting them away from the cytosol [47]. Even all *VrHMA* genes, except *VrHMA3*, were upregulated in both roots and leaves in response to Zn^2+^ stress, either gradually with increment of time upto 72 hours or after a specific duration (24 hours or 48 hours) of treatment, indicating their putative role in zinc influx to cytoplasm and efflux to vacuoles as reported in *A. thaliana*, *O. sativa* and *A. catechu* [31, 73]. A comparable expression pattern was evident in case of *VrHMA1* and *VrHMA5* in both root and leaf tissues. Similarly, the high *VrHMA7* root-specific expression can be corelated with *A*. *halleri*, *AhHMA4* where its high expression in roots might be mediating in the higher rates of root to shoot zinc translocation, further helping in the zinc homoeostasis process in root tissues [51]. In leaves, some of the genes, such as *VrHMA4, VrHMA6* and *VrHMA7* were upregulated after specific duration of zinc stress and thereafter downregulated, which could be attributed their role on sequestration of increased accumulation of zinc in side cytosol [14]. In response to Cd^2+^ stress, eight out of nine *VrHMA* genes were overexpressed both in roots (except *VrHMA1*) and leaves (except *VrHMA5*) either gradually with increment of time or after a specific duration (up to 72 hours) of treatment. Alike Zn^2+^ response, *VrHMA5* and *VrHMA7,* belong to Zn/Co/Cd/Pb-ATPases subgroup, showed incremental upregulation in roots in response to Cd^2+^ stress, whereas *VrHMA1* not expressed even after 72 hours of Cd^2+^ stress treatment. This indicates their putative role in systemic redistribution and sequestration of Cd [15]. The expression of *VrHMA* genes (2, 6, 8, 9) were upregulated in roots after 24 hours, and then decreases gradually with time in roots, which can be attributed to the negative regulation of expression of these genes under Cd^2+^ stress [25, 82]. Alike roots, *VrHMA4*, *VrHMA7* and *VrHMA8* elicited in its expression in leaves after 24 hours of induction, and then subsided upto 72 hours of the treatment, and this can be attributed to the Cd toxicity in leaves [82]. In response to Cu^2+^ stress (50µM CuSO_4_), five out of nine *VrHMA* genes (*VrHMA-1, -2, -4, -6, -7* and *-8*) were overexpressed in roots after 24 hours of treatment and then decreased gradually with increment of time, which could be attributed to increment of Cu ions toxicity as reported in *M*. *truncatula* [44]. In contrast, five out of nine *VrHMA* genes (*VrHMA-1, -2, -3, -8* and -*9*) showed incremental upregulated expression in leaves upto 72 hours of treatment similar to *P. trichocarpa*, which might be due to the involvement of these *VrHMAs* in the transport and detoxification through sequestration of Cu in leaves as half of the Cu^2+^ stores chloroplast [38, 59, 78]. In particular, the chloroplast localized *VrHMAs* (1 and 9) are likely to be associated with transfer Cu^2+^ to Cu-Zn superoxide dismutase and plastocyanin as reported in *A. thaliana* and *M. truncatula* [1, 44]. Interestingly, the expression levels of *VrHMA5*, *VrHMA6* and *VrHMA1* in the roots after 48h and 72 h of Cu stress were 18, 12 and 4.6-fold upregulated in comparison to control. Similarly, after 48h and 72h of Cu stress, the expression levels of *VrHMA1*, *VrHMA2* and *VrHMA8* in the leaves were 29, 13 and 7-fold increase in comparison to control. This copper stress induced differential expression pattern in *V*. *radiata* could be attributed to the role of VrHMAs in the initiation of Cu^2+^ uptake by roots and its transportation to vascular tissues of the stems and leaves, and it also indicated that leaves might be the principal plant part of Cu detoxification, whereas roots mediate uptake and inhibition Cu as reported *M. truncatula* [39, 44].

## Conclusions

HMA transporters play a crucial role in the uptake, distribution and sequestration of essential, nonessential and even toxic metals ions in plants. In this study, nine *VrHMA* genes were identified in the Mung bean genome, which were broadly grouped to Zn/Co/Cd/Pb-ATPases and Cu/Ag-ATPases along with their homologs from *A. thaliana*, *M. truncatula* and *G. max*. These *VrHMA* genes had conserved and divergent gene structure, protein architecture, motif distribution pattern. These genes could be used for functional genomic studies pertaining to Zn biofortification, modulation of heavy metal stress, mechanisms of detoxification and their role in plant development under abiotic stress. The expression profiles of *VrHMA* genes in root and leaf tissues in response to heavy metal (Zn, Cd and Cu) stress indicate their role in the uptake, transport and sequestration alike their homologs from model dicots as well as model legumes. Interestingly *VrHMA5*, belongs to Zn/Co/Cd/Pb-ATPases subgroup, showed incremental upregulation in roots under all three heavy metal stress, whereas it only showed upregulated expression in the leaf tissues under Zn stress, which indicates its role in vascular transport in *V. radiata*. The findings of the present study are valuable for functional genomic exploration of the molecular mechanisms responding to atleast heavy metal stress vis-à-vis Zn homeostasis in *V. radiata*.

## Abbreviations

ABRE: ABA responsive elements
ARE: Auxin responsive element
CDS: Coding sequences
CAE: Cis-acting elements
ERE: Ethylene responsive element
GRAVY: grand average of hydropathicity
HMA: Heavy metal ATPase
HMM: Hidden Markov Model
LIS: Legume Information System
LRE: Light regulatory element
LTRE: Low temperature responsive element
MBD: Metal binding domains
MUSCLE: multiple sequence comparison by log-expectation
pI: Isoelectric point
PLH: Pore Line helices
qRT-PCR: Quantitative Real-time Polymerase Chain Reaction
TMH: Trans-membrane Helices

---

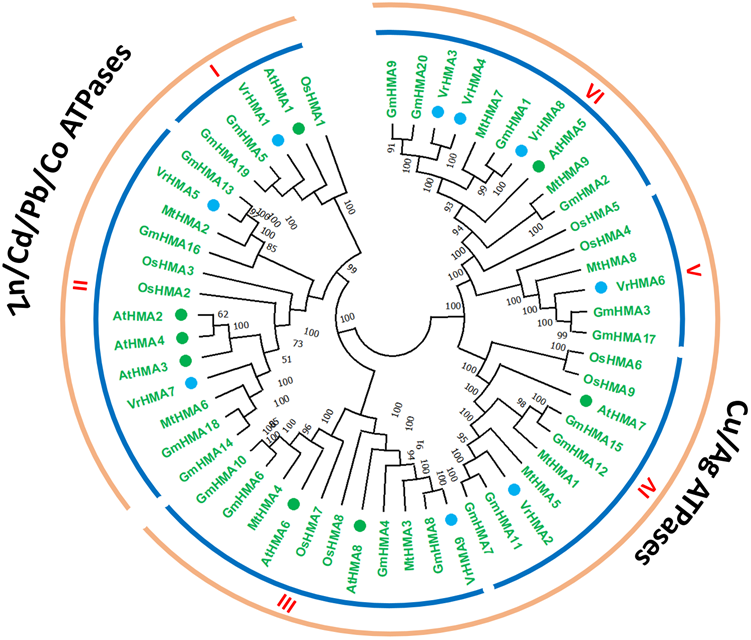

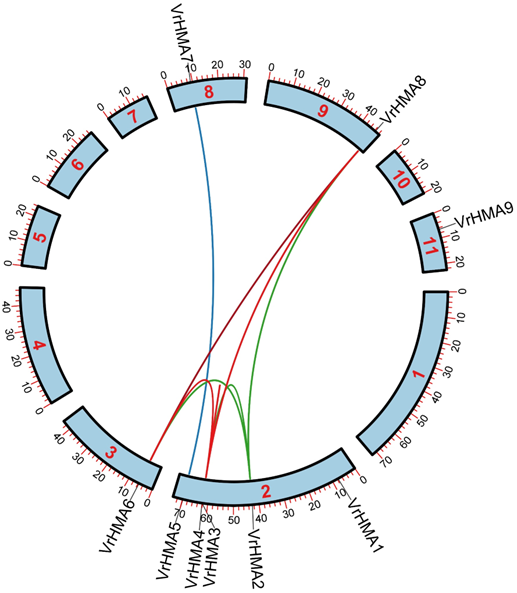

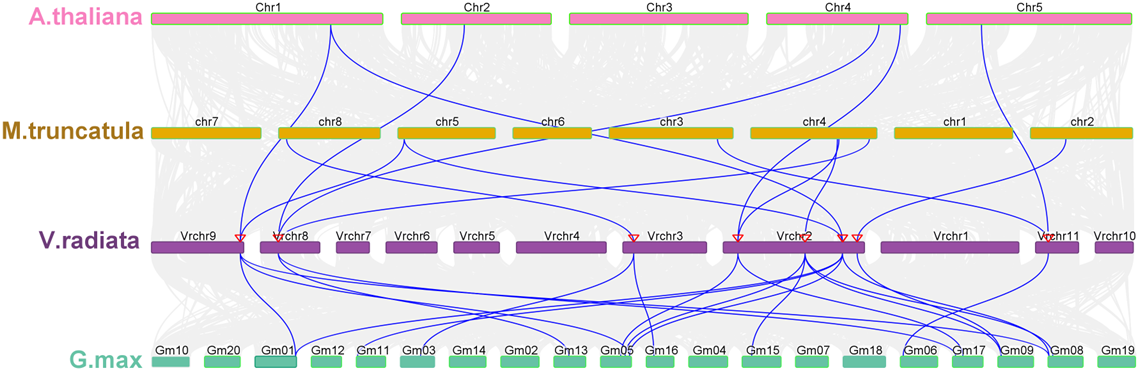

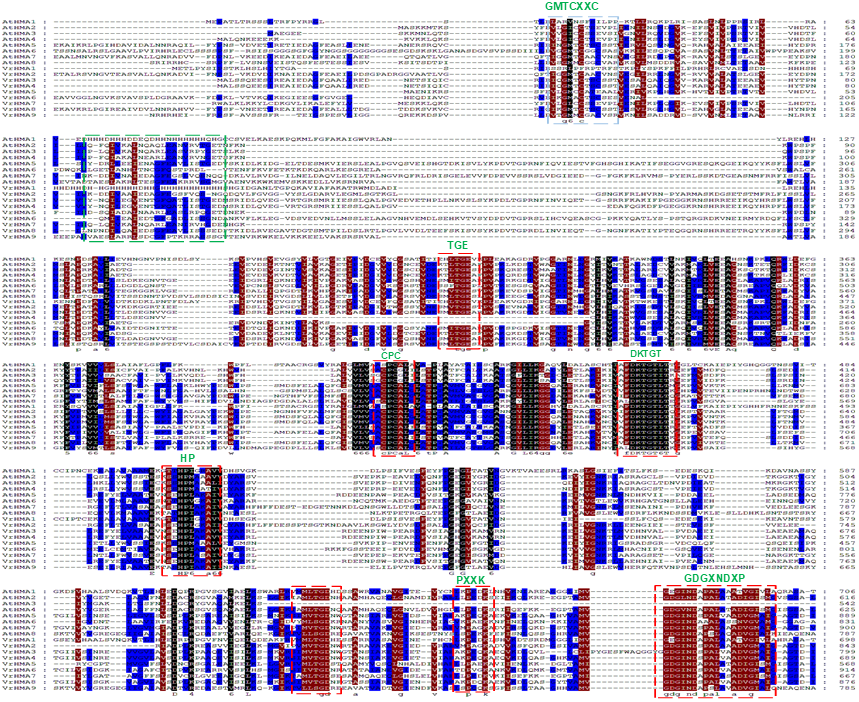

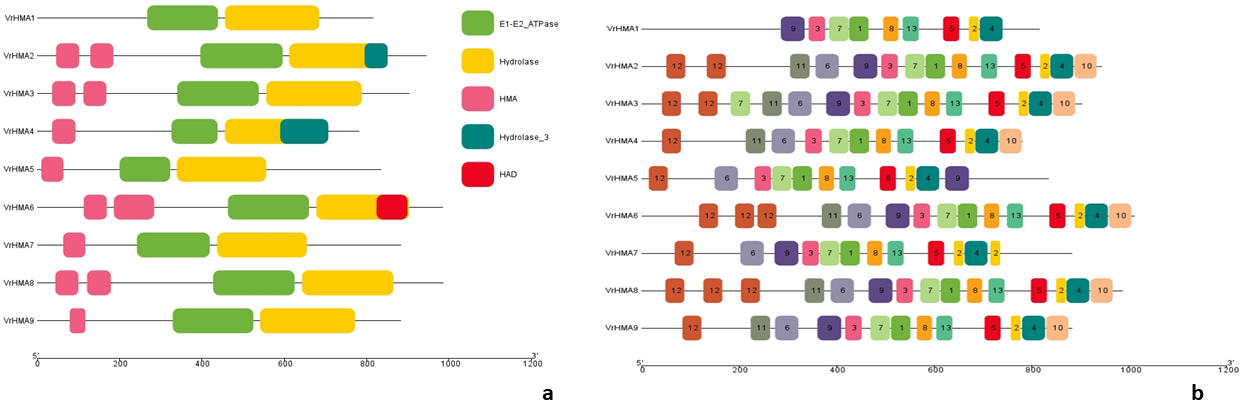

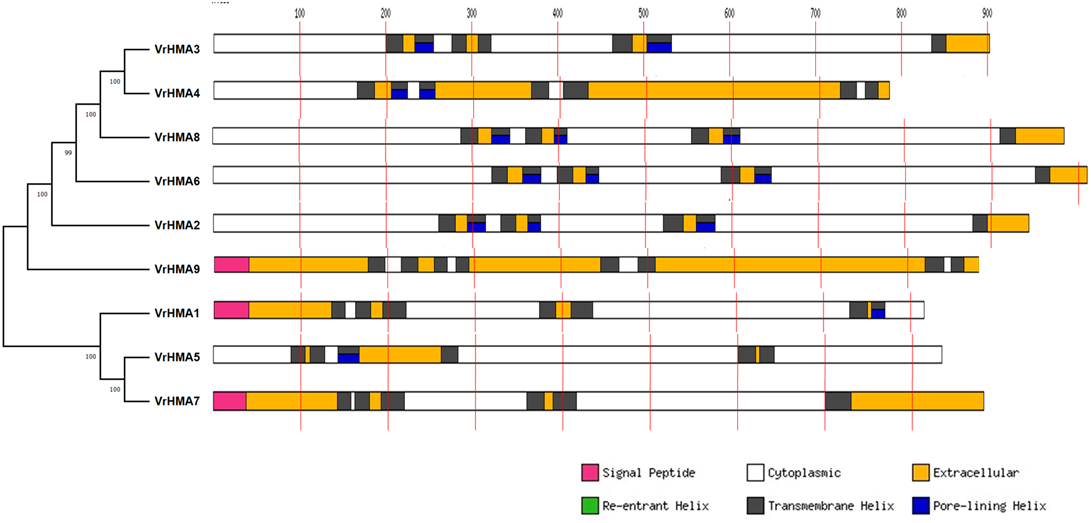

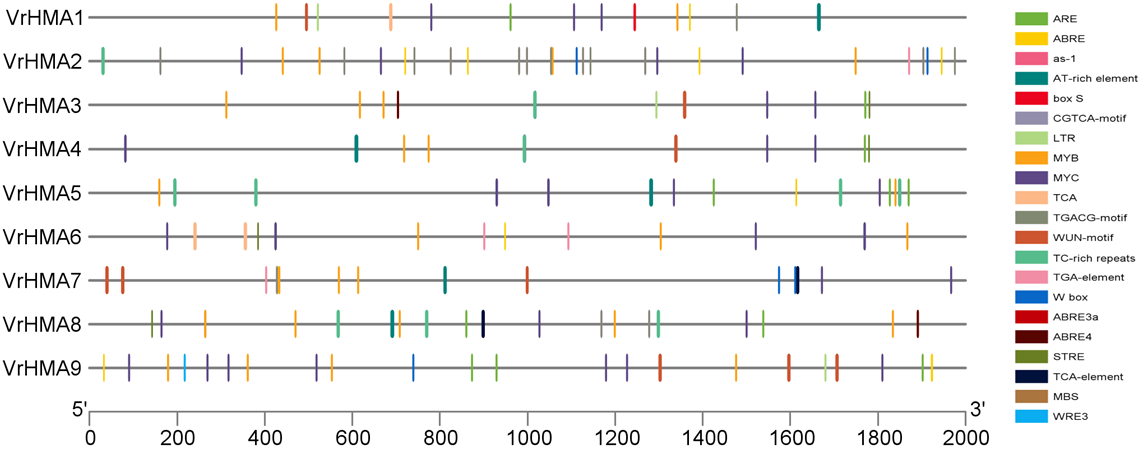

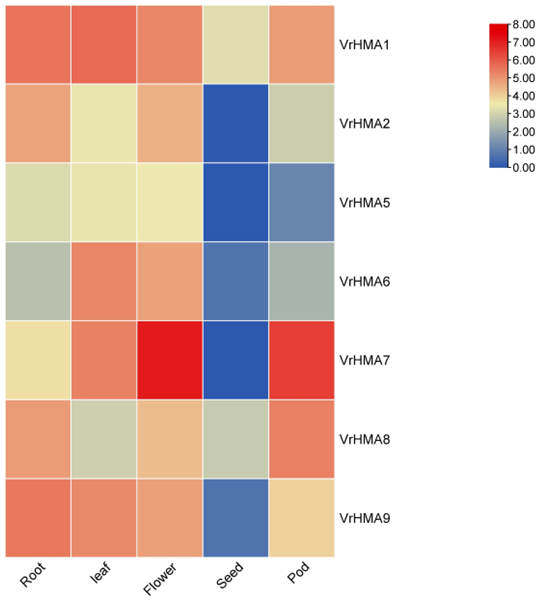

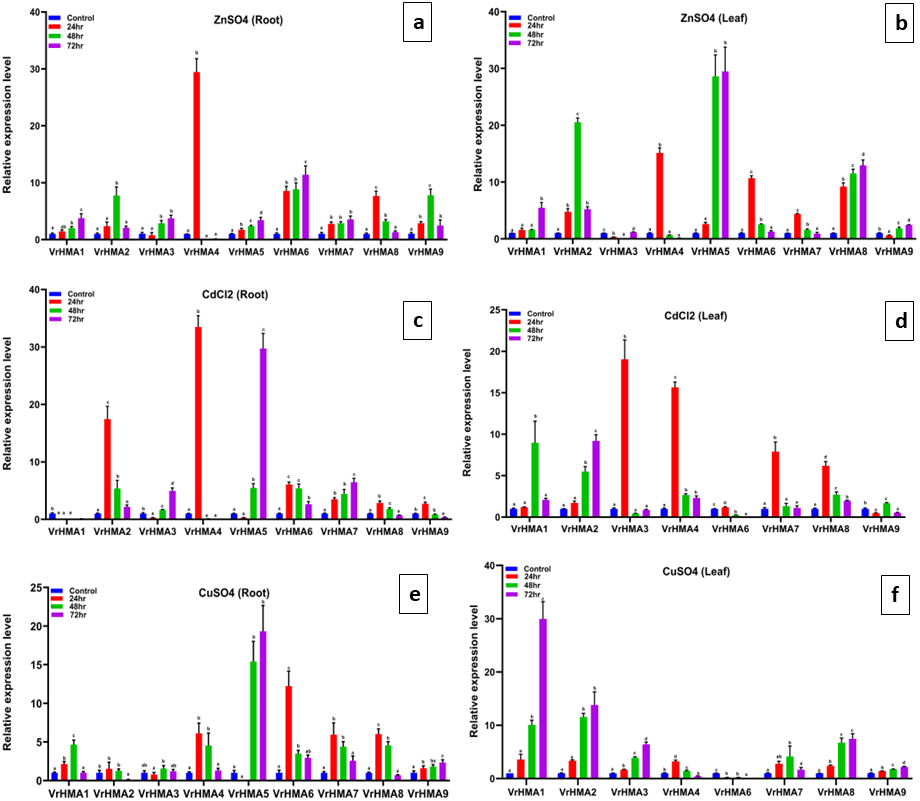

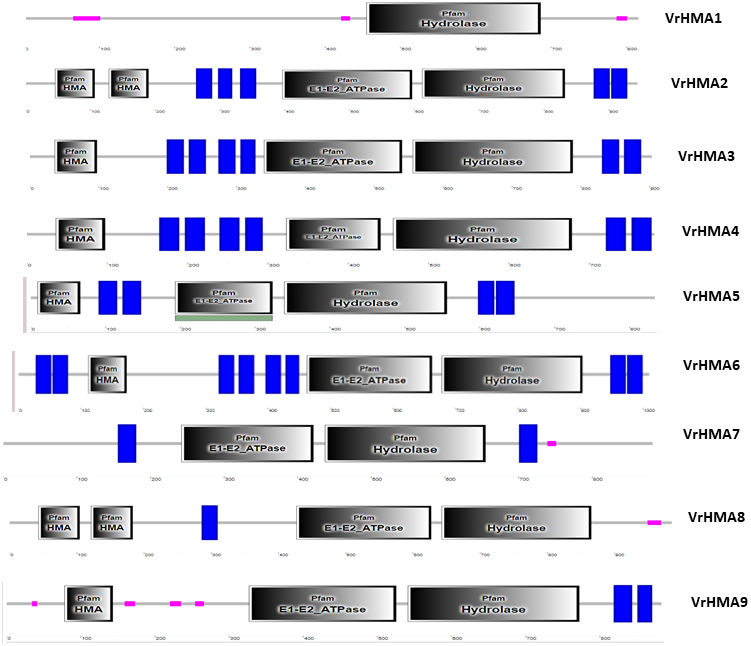

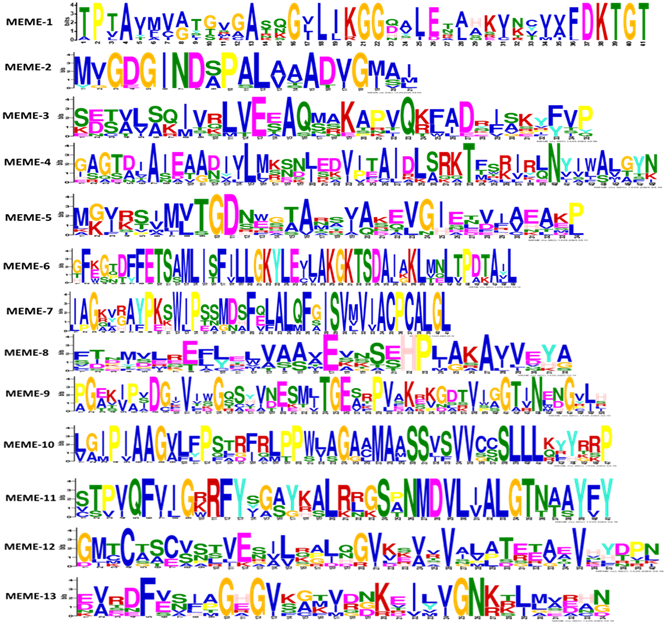

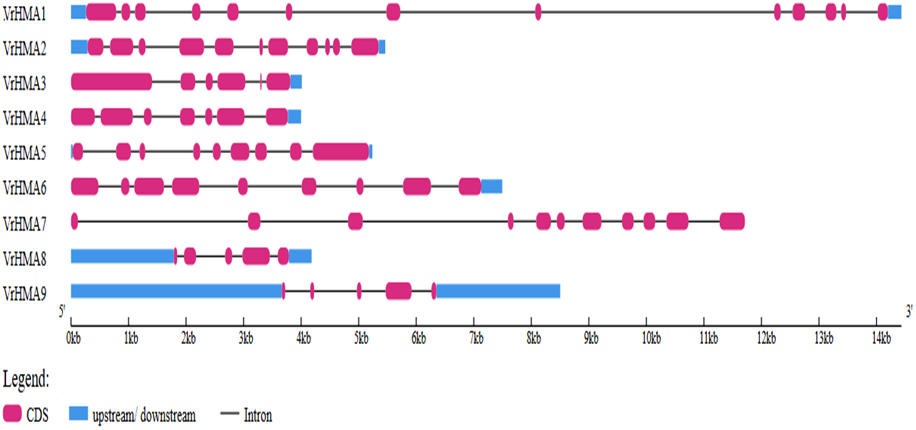

**Supplementary Table 1:**
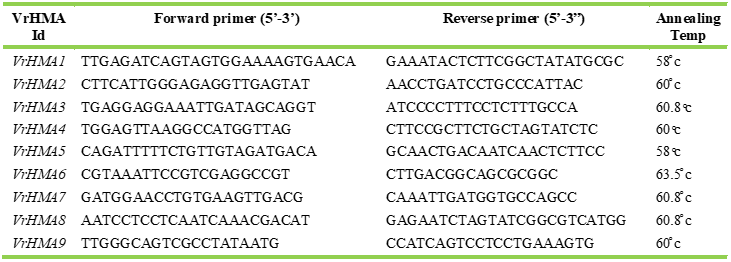
List of primers used for quantitative real time(qRT) PCR analysis of VrHMA genes.

**Supplementary Table 2:**
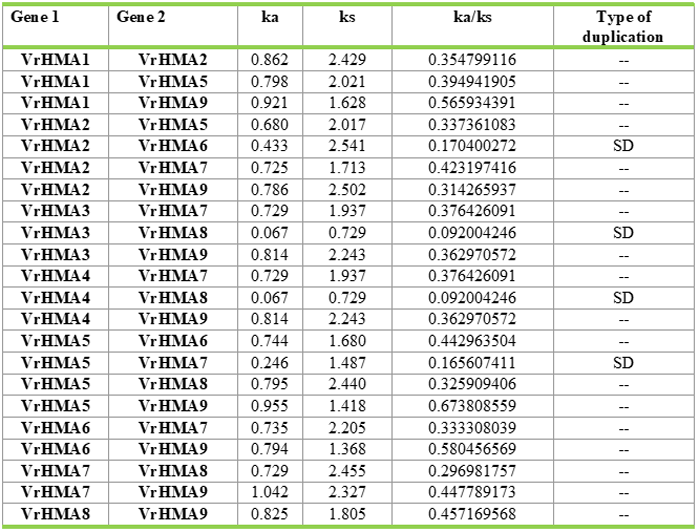
Estimation of codon-based (Synonymous vs Non-synonymous) evolutionary divergence between VrHMA sequences determined using Mega-X software.

**Supplementary Table 3.**
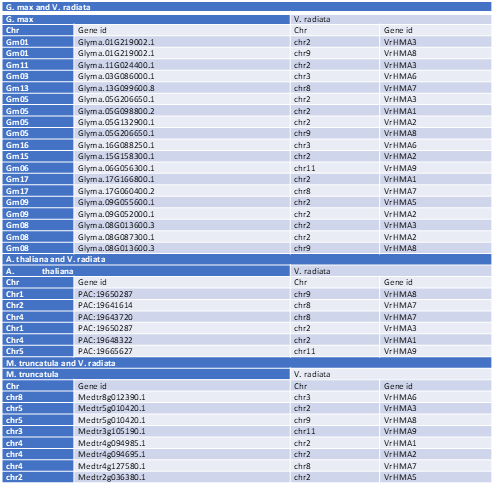
Summary of interspecifc synteny analyses among HMA gene family *V. radiata, A. thaliana, G. max* and *M. truncatula* showing homologous gene pairs.

**Supplementary Table 4:**
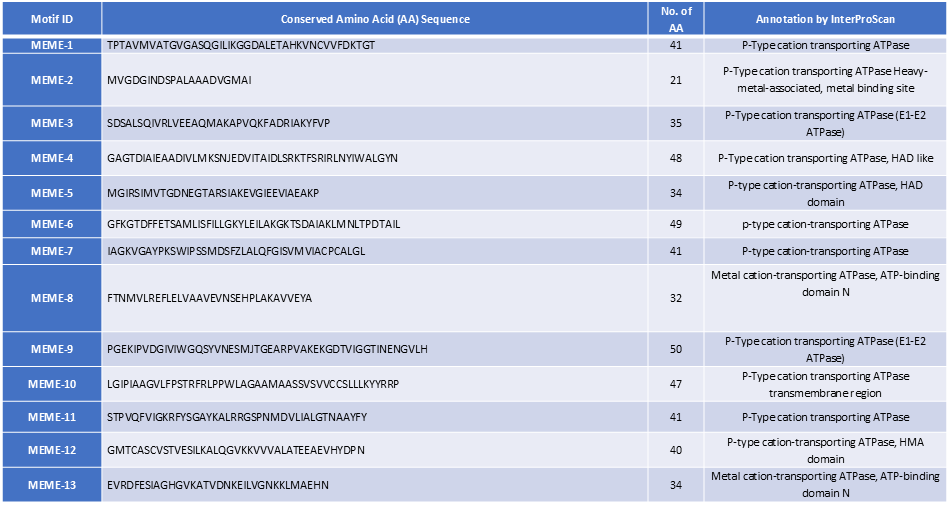
Thirteen conserved motifs present in nine *VrHMA* genes of Mung bean deduced by MEME suite and their putative functions as annotated by lnterProScan.

**Supplementary Table 5:**
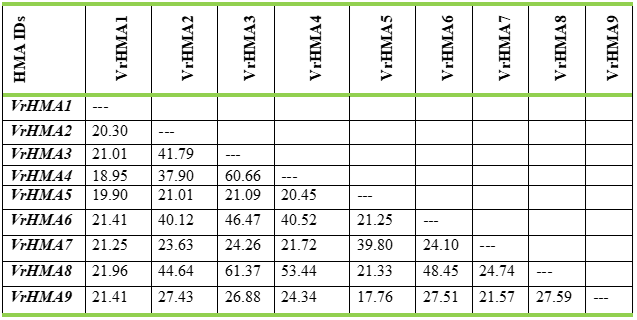
Amino acid sequence similarity(%) among the nine VrHMA proteins of Mung bean.

**Supplementary Table 6:**
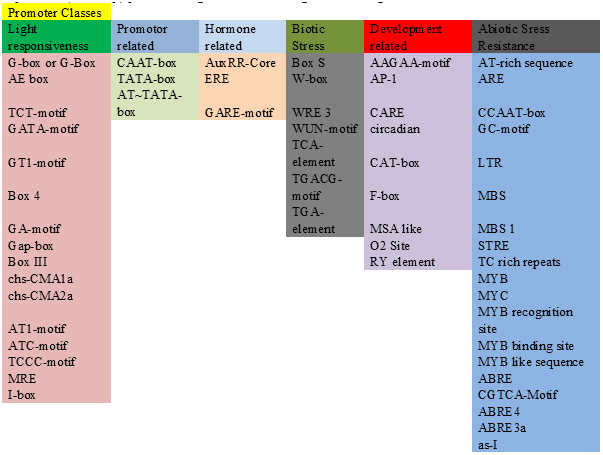
Identification of putative transcription regulatory cis-acting elements in the upstream (-2000bp) promoter region of 9VrHMA genes in Mung bean.

**Supplementary Table 7:**
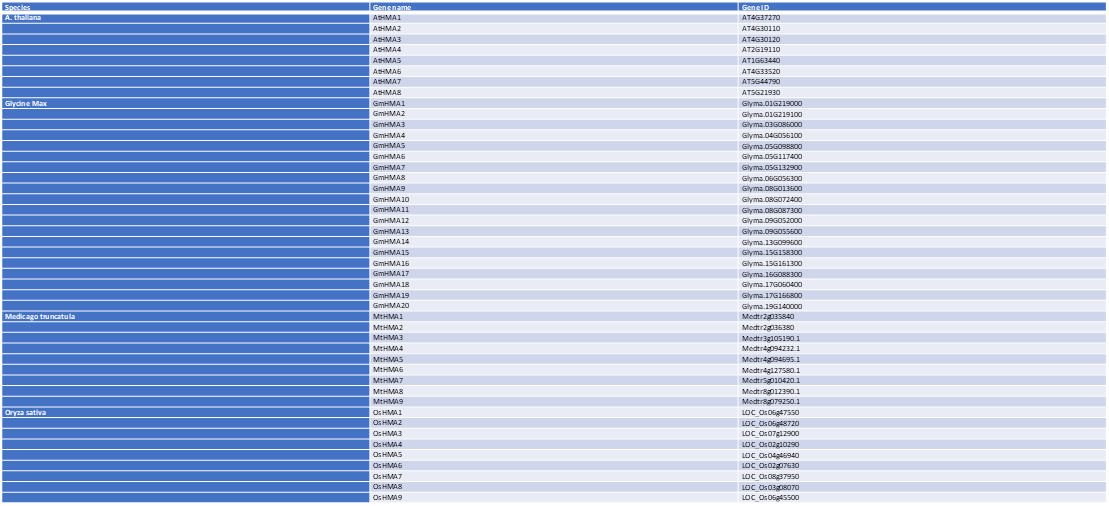
HMA genes of model plants *A. thaliana* and *O. sativa,* and two model legume species *G. max* and *M. truncatula* used for interspecific phylogenetic studies.

## References

1. Abdel-Ghany SE, Müller-Moulé P, Niyogi KK, Pilon M, Shikanai T. Two P-Type ATPases Are Required for Copper Delivery in *Arabidopsis thaliana* Chloroplasts. Plant Cell. 2005;17:1233–51. 10.1105/tpc.104.030452.

2. AndrésLColás N, Sancenón V, RodríguezLNavarro S, Mayo S, Thiele DJ, Ecker JR, et al. The Arabidopsis heavy metal PLtype ATPase HMA5 interacts with metallochaperones and functions in copper detoxification of roots. Plant J. 2006;45:225–36. 10.1111/j.1365-313X.2005.02601.x.

3. Bailey TL, Boden M, Buske FA, Frith M, Grant CE, Clementi L, et al. MEME SUITE: tools for motif discovery and searching. Nucleic Acids Res. 2009;37:W202–8. 10.1093/nar/gkp335.

4. Batool TS, Aslam R, Gul A, Paracha RZ, Ilyas M, De Abreu K, et al. Genome-wide analysis of heavy metal ATPases (HMAs) in Poaceae species and their potential role against copper stress in Triticum aestivum. Sci Rep. 2023;13:7551. 10.1038/s41598-023-32023-7.

5. Boutigny S, Sautron E, Finazzi G, Rivasseau C, Frelet-Barrand A, Pilon M, et al. HMA1 and PAA1, two chloroplast-envelope PIB-ATPases, play distinct roles in chloroplast copper homeostasis. J Exp Bot. 2014;65:1529–40. 10.1093/jxb/eru020.

6. Cai H, Huang S, Che J, Yamaji N, Ma JF. The tonoplast-localized transporter OsHMA3 plays an important role in maintaining Zn homeostasis in rice. J Exp Bot. 2019;70:2717–25. 10.1093/jxb/erz091.

7. Capdevila M, Atrian S. Metallothionein protein evolution: a miniassay. JBIC J Biol Inorg Chem. 2011;16:977–89. 10.1007/s00775-011-0798-3.

8. Chalhoub B, Denoeud F, Liu S, Parkin IAP, Tang H, Wang X, et al. Early allopolyploid evolution in the post-Neolithic *Brassica napus* oilseed genome. Science. 2014;345:950–3. 10.1126/science.1253435.

9. Chen C, Wu Y, Li J, Wang X, Zeng Z, Xu J, et al. TBtools-II: A “one for all, all for one” bioinformatics platform for biological big-data mining. Mol Plant. 2023;16:1733–42. 10.1016/j.molp.2023.09.010.

10. Cobbett CS, Hussain D, Haydon MJ. Structural and functional relationships between type 1_B_ heavy metalLtransporting PLtype ATPases in *Arabidopsis*. New Phytol. 2003;159:315–21. 10.1046/j.1469-8137.2003.00785.x.

11. De Castro E, Sigrist CJA, Gattiker A, Bulliard V, Langendijk-Genevaux PS, Gasteiger E, et al. ScanProsite: detection of PROSITE signature matches and ProRule-associated functional and structural residues in proteins. Nucleic Acids Res. 2006;34:W362–5. 10.1093/nar/gkl124.

12. Deng F, Yamaji N, Xia J, Ma JF. A Member of the Heavy Metal P-Type ATPase OsHMA5 Is Involved in Xylem Loading of Copper in Rice. Plant Physiol. 2013;163:1353–62. 10.1104/pp.113.226225.

13. D’Ovidio R, Raiola A, Capodicasa C, Devoto A, Pontiggia D, Roberti S, et al. Characterization of the Complex Locus of Bean Encoding Polygalacturonase-Inhibiting Proteins Reveals Subfunctionalization for Defense against Fungi and Insects. Plant Physiol. 2004;135:2424–35. 10.1104/pp.104.044644.

14. Eren E, Argüello JM. Arabidopsis HMA2, a divalent heavy metal-transporting P(IB)-type ATPase, is involved in cytoplasmic Zn2+ homeostasis. Plant Physiol. 2004;136:3712–23. 10.1104/pp.104.046292.

15. Fang X, Wang L, Deng X, Wang P, Ma Q, Nian H, et al. Genome-wide characterization of soybean P 1B -ATPases gene family provides functional implications in cadmium responses. BMC Genomics. 2016; 17:376. 10.1186/s12864-016-2730-2.

16. Furukawa Y, Lim C, Tosha T, Yoshida K, Hagai T, Akiyama S, et al. Identification of a novel zinc-binding protein, C1orf123, as an interactor with a heavy metal-associated domain. Dmitriev OY, editor. PLOS ONE. 2018;13:e0204355. 10.1371/journal.pone.0204355.

17. Gasteiger E, Gattiker A, Hoogland C, Ivanyi I, Appel RD, Bairoch A. ExPASy: The proteomics server for in-depth protein knowledge and analysis. Nucleic Acids Res. 2003;31:3784–8. 10.1093/nar/gkg563.

18. Guo Z, Orädd F, Bågenholm V, Grønberg C, Ma JF, Ott P, et al. Diverse roles of the metal binding domains and transport mechanism of copper transporting P-type ATPases. Nat Commun. 2024;15:2690. 10.1038/s41467-024-47001-4.

19. Ha J, Satyawan D, Jeong H, Lee E, Cho K, Kim MY, et al. A nearLcomplete genome sequence of mungbean ( *Vigna radiata* L.) provides key insights into the modern breeding program. Plant Genome. 2021;14:e20121. 10.1002/tpg2.20121.

20. Hambidge M. Human Zinc Deficiency. J Nutr. 2000;130:1344S–1349S. 10.1093/jn/130.5.1344S.

21. He G, Qin L, Tian W, Meng L, He T, Zhao D. Heavy Metal Transporters-Associated Proteins in Solanum tuberosum: Genome-Wide Identification, Comprehensive Gene Feature, Evolution and Expression Analysis. Genes. 2020;11:1269. 10.3390/genes11111269.

22. Hirayama T, Kieber JJ, Hirayama N, Kogan M, Guzman P, Nourizadeh S, et al. RESPONSIVE-TO-ANTAGONIST1, a Menkes/Wilson Disease–Related Copper Transporter, Is Required for Ethylene Signaling in Arabidopsis. Cell. 1999;97:383–93. 10.1016/S0092-8674(00)80747-3.

23. Hou D, Yousaf L, Xue Y, Hu J, Wu J, Hu X, et al. Mung Bean (Vigna radiata L.): Bioactive Polyphenols, Polysaccharides, Peptides, and Health Benefits. Nutrients. 2019;11:1238. 10.3390/nu11061238.

24. Hu B, Jin J, Guo A-Y, Zhang H, Luo J, Gao G. GSDS 2.0: an upgraded gene feature visualization server. Bioinformatics. 2015;31:1296–7. 10.1093/bioinformatics/btu817.

25. Huang Q, Qiu W, Yu M, Li S, Lu Z, Zhu Y, et al. Genome-Wide Characterization of Sedum plumbizincicola HMA Gene Family Provides Functional Implications in Cadmium Response. Plants. 2022;11:215. 10.3390/plants11020215.

26. Huang Y, Niu B, Gao Y, Fu L, Li W. CD-HIT Suite: a web server for clustering and comparing biological sequences. Bioinformatics. 2010;26:680–2. 10.1093/bioinformatics/btq003.

27. Hussain D, Haydon MJ, Wang Y, Wong E, Sherson SM, Young J, et al. P-Type ATPase Heavy Metal Transporters with Roles in Essential Zinc Homeostasis in Arabidopsis. Plant Cell. 2004;16:1327–39. 10.1105/tpc.020487.

28. Jamsheer K M, Kumar M. Transcription Factors as Zinc Sensors in Plants. Trends Plant Sci. 2021;26:761–3. 10.1016/j.tplants.2021.04.008..

29. Jasrotia RS, Jaiswal S, Yadav PK, Raza M, Iquebal MA, Rai A, et al. Genome-Wide Analysis of HSP70 Family Protein in *Vigna radiata* and Coexpression Analysis Under Abiotic and Biotic Stress. J Comput Biol. 2020;27:738–54. 10.1089/cmb.2019.0166.

30. Kang YJ, Kim SK, Kim MY, Lestari P, Kim KH, Ha B-K, et al. Genome sequence of mungbean and insights into evolution within Vigna species. Nat Commun. 2014;5:5443. 10.1038/ncomms6443.

31. Khan NM, Ali A, Wan Y, Zhou G. Genome-wide identification of heavy-metal ATPases genes in Areca catechu: investigating their functionality under heavy metal exposure. BMC Plant Biol. 2024;24:484. 10.1186/s12870-024-05201-6.

32. Kim Y, Choi H, Segami S, Cho H, Martinoia E, Maeshima M, et al. AtHMA1 contributes to the detoxification of excess Zn(II) in Arabidopsis. Plant J. 2009;58:737–53. 10.1111/j.1365-313X.2009.03818.x.

33. Krogh A, Larsson B, Von Heijne G, Sonnhammer ELL. Predicting transmembrane protein topology with a hidden markov model: application to complete genomes11Edited by F. Cohen. J Mol Biol. 2001;305:567–80. 10.1006/jmbi.2000.4315.

34. Kumar S, Stecher G, Li M, Knyaz C, Tamura K. MEGA X: Molecular Evolutionary Genetics Analysis across Computing Platforms. Battistuzzi FU, editor. Mol Biol Evol. 2018;35:1547–9. 10.1093/molbev/msy096.

35. Lee S, Kim Y-Y, Lee Y, An G. Rice P1B-Type Heavy-Metal ATPase, OsHMA9, Is a Metal Efflux Protein. Plant Physiol. 2007;145:831–42. 10.1104/pp.107.102236.

36. Lescot M. PlantCARE, a database of plant cis-acting regulatory elements and a portal to tools for in silico analysis of promoter sequences. Nucleic Acids Res. 2002;30:325–7. 10.1093/nar/30.1.325.

37. Letunic I, Doerks T, Bork P. SMART 7: recent updates to the protein domain annotation resource. Nucleic Acids Res. 2012;40:D302–5. 10.1093/nar/gkr9.

38. Li D, Xu X, Hu X, Liu Q, Wang Z, Zhang H, et al. Genome-Wide Analysis and Heavy Metal-Induced Expression Profiling of the HMA Gene Family in Populus trichocarpa. Front Plant Sci. 2015;6. 10.3389/fpls.2015.01149.

39. Li J, Zhang Z, Shi G. Genome-Wide Identification and Expression Profiling of Heavy Metal ATPase (HMA) Genes in Peanut: Potential Roles in Heavy Metal Transport. Int J Mol Sci. 2024;25:613. 10.3390/ijms25010613.

40. Li N, Xiao H, Sun J, Wang S, Wang J, Chang P, et al. Genome-wide analysis and expression profiling of the HMA gene family in Brassica napus under cd stress. Plant Soil. 2018;426:365–81. 10.1007/s11104-018-3637-2.

41. Likhith RK, Alagarasan G, Muthurajan R, Parasuraman B, Subramanian R. Genome wide identification of mungbean (Vigna radiata [L.] R. Wilczek) Late Embryogenesis Abundant (LEA) protein gene family. Isr J Plant Sci. 2021;69:79–86. 10.1163/22238980-bja10049.

42. Liu C, Wang Y, Peng J, Fan B, Xu D, Wu J, et al. High-quality genome assembly and pan-genome studies facilitate genetic discovery in mung bean and its improvement. Plant Commun. 2022;3:100352. 10.1016/j.xplc.2022.100352.

43. Livak KJ, Schmittgen TD. Analysis of Relative Gene Expression Data Using Real-Time Quantitative PCR and the 2−ΔΔCT Method. Methods. 2001;25:402–8. 10.1006/meth.2001.1262.

44. Ma Y, Wei N, Wang Q, Liu Z, Liu W. Genome-wide identification and characterization of the heavy metal ATPase (HMA) gene family in Medicago truncatula under copper stress. Int J Biol Macromol. 2021; 193:893–902. 10.1016/j.ijbiomac.2021.10.197.

45. Mehta N, Rao P, Saini R. EXPLORATION OF THE ANTIBACTERIAL, ANTIOXIDANT AND ANTICANCER POTENTIAL OF THE SEED COAT EXTRACT OF MUNGBEAN (VIGNA RADIATA L. WILCZEK). PLANT Arch. 2021;21. 10.51470/PLANTARCHIVES.2021.v21.no1.222.

46. Mills RF, Francini A, Ferreira Da Rocha PSC, Baccarini PJ, Aylett M, Krijger GC, et al. The plant P_1B_ Ltype ATPase AtHMA4 transports Zn and Cd and plays a role in detoxification of transition metals supplied at elevated levels. FEBS Lett. 2005;579:783–91. 10.1016/j.febslet.2004.12.040.

47. Morel M, Crouzet J, Gravot A, Auroy P, Leonhardt N, Vavasseur A, et al. AtHMA3, a P1B-ATPase Allowing Cd/Zn/Co/Pb Vacuolar Storage in Arabidopsis. Plant Physiol. 2009;149:894–904. 10.1104/pp.108.130294.

48. Nag A, Gupta K, Dubey N, Mishra SK, Panigrahi J. Genomic characterization of ZIP genes in pigeonpea (CcZIP) and their expression analysis among the genotypes with contrasting host response to pod borer. Physiol Mol Biol Plants. 2021;27:2787–804. 10.1007/s12298-021-01111-1.

49. Nazmul Hasan Md, Islam S, Bhuiyan FH, Arefin S, Hoque H, Azad Jewel N, et al. Genome wide analysis of the heavy-metal-associated (HMA) gene family in tomato and expression profiles under different stresses. Gene. 2022;835:146664. 10.1016/j.gene.2022.146664.

50. Nicholas KB, Nicholas HB. GeneDoc: a tool for editing and annotating multiple sequence alignments. 1997. https://api.semanticscholar.org/CorpusID:81058551.

51. Nouet C, Charlier J-B, Carnol M, Bosman B, Farnir F, Motte P, et al. Functional analysis of the three *HMA4* copies of the metal hyperaccumulator *Arabidopsis halleri*. J Exp Bot. 2015;66:5783–95. 10.1093/jxb/erv280.

52. Nugent T, Ward S, Jones DT. The MEMPACK alpha-helical transmembrane protein structure prediction server. Bioinformatics. 2011;27:1438–9. 10.1093/bioinformatics/btr096.

53. Olsen LI, Palmgren MG. Many rivers to cross: the journey of zinc from soil to seed. Front Plant Sci. 2014;5. 10.3389/fpls.2014.00030.

54. Petersen TN, Brunak S, Von Heijne G, Nielsen H. SignalP 4.0: discriminating signal peptides from transmembrane regions. Nat Methods. 2011;8:785–6. 10.1038/nmeth.1701.

55. Pita-Barbosa A, Ricachenevsky FK, Wilson M, Dottorini T, Salt DE. Transcriptional plasticity buffers genetic variation in zinc homeostasis. Sci Rep. 2019;9:19482. 10.1038/s41598-019-55736-0.

56. Quevillon E, Silventoinen V, Pillai S, Harte N, Mulder N, Apweiler R, et al. InterProScan: protein domains identifier. Nucleic Acids Res. 2005;33:W116–20. 10.1093/nar/gki442.

57. Santner A, Estelle M. Recent advances and emerging trends in plant hormone signalling. Nature. 2009; 459:1071–8. 10.1038/nature08122.

58. Satoh-Nagasawa N, Mori M, Nakazawa N, Kawamoto T, Nagato Y, Sakurai K, et al. Mutations in Rice (Oryza sativa) Heavy Metal ATPase 2 (OsHMA2) Restrict the Translocation of Zinc and Cadmium. Plant Cell Physiol. 2012;53:213–24. 10.1093/pcp/pcr166.

59. Sautron E, Mayerhofer H, Giustini C, Pro D, Crouzy S, Ravaud S, et al. HMA6 and HMA8 are two chloroplast Cu+-ATPases with different enzymatic properties. Biosci Rep. 2015;35:e00201. 10.1042/BSR20150065.

60. Seigneurin-Berny D, Gravot A, Auroy P, Mazard C, Kraut A, Finazzi G, et al. HMA1, a New Cu-ATPase of the Chloro plast Envelope, Is Essential for Growth under Adverse Light Conditions. J Biol Chem . 2006;281:2882–92. 10.1074/jbc.M508333200.

61. Shahmuradov IA, Solovyev VV. Nsite, NsiteH and NsiteM computer tools for studying transcription regulatory elements. Bioinformatics. 2015;31:3544–5. 10.1093/bioinformatics/btv404.

62. Singh A, Gupta M, Kumar S, Chawla H, Mehra M, Kumar K, et al. Posttranslational modifications and metal stress tolerance in plants. Biostimulants Alleviation Met Toxic Plants. Elsevier; 2023. p. 511–31. 10.1016/B978-0-323-99600-6.00001-3.

63. Singh S, Parihar P, Singh R, Singh VP, Prasad SM. Heavy Metal Tolerance in Plants: Role of Transcriptomics, Proteomics, Metabolomics, and Ionomics. Front Plant Sci. 2016;6. 10.3389/fpls.2015.01143.

64. Smith AT, Smith KP, Rosenzweig AC. Diversity of the metal-transporting P1B-type ATPases. JBIC J Biol Inorg Chem. 2014;19:947–60. 10.1007/s00775-014-1129-2.

65. Tabata K, Kashiwagi S, Mori H, Ueguchi C, Mizuno T. Cloning of a cDNA encoding a putative metal-transporting P-type ATPase from Arabidopsis thaliana. Biochim Biophys Acta BBA - Biomembr. 1997; 1326:1–6. 10.1016/S0005-2736(97)00064-3.

66. Takahashi R, Bashir K, Ishimaru Y, Nishizawa NK, Nakanishi H. The role of heavy-metal ATPases, HMAs, in zinc and cadmium transport in rice. Plant Signal Behav. 2012;7:1605–7. 10.4161/psb.22454.

67. Takahashi R, Ishimaru Y, Shimo H, Ogo Y, Senoura T, Nishizawa NK, et al. The OsHMA2 transporter is involved in rootLtoLshoot translocation of Zn and Cd in rice. Plant Cell Environ. 2012;35:1948–57. 10.1111/j.1365-3040.2012.02527.x.

68. Talakayala A, Mekala GK, Reddy MK, Ankanagari S, Garladinne M. Manipulating resistance to mungbean yellow mosaic virus in greengram (Vigna radiata L): Through CRISPR/Cas9 mediated editing of the viral genome. Front Sustain Food Syst. 2022;6:911574. 10.3389/fsufs.2022.911574.

69. Tang H, Krishnakumar V, Bidwell S, Rosen B, Chan A, Zhou S, et al. An improved genome release (version Mt4.0) for the model legume Medicago truncatula.

70. The Arabidopsis Genome Initiative. Analysis of the genome sequence of the flowering plant Arabidopsis thaliana. Nature. 2000;408:796–815. 10.1038/35048692.

71. Thingholm TE, Rönnstrand L, Rosenberg PA. Why and how to investigate the role of protein phosphorylation in ZIP and ZnT zinc transporter activity and regulation. Cell Mol Life Sci. 2020;77:3085–102. 10.1007/s00018-020-03473-3.

72. Traverso JA, Meinnel T, Giglione C. Expanded impact of protein N-myristoylation in plants. Plant Signal Behav. 2008;3:501–2. 10.4161/psb.3.7.6039.

73. Verret F, Gravot A, Auroy P, Leonhardt N, David P, Nussaume L, et al. Overexpression of AtHMA4 enhances rootLtoLshoot translocation of zinc and cadmium and plant metal tolerance. FEBS Lett. 2004;576:306–12. 10.1016/j.febslet.2004.09.023.

74. Verret F, Gravot A, Auroy P, Preveral S, Forestier C, Vavasseur A, et al. Heavy metal transport by AtHMA4 involves the NLterminal degenerated metal binding domain and the CLterminal His_11_ stretch. FEBS Lett. 2005;579:1515–22. 10.1016/j.febslet.2005.01.065.

75. Williams LE, Mills RF. P1B-ATPases – an ancient family of transition metal pumps with diverse functions in plants. Trends Plant Sci. 2005;10:491–502. 10.1016/j.tplants.2005.08.008.

76. Wong CKE, Cobbett CS. HMA PLtype ATPases are the major mechanism for rootLtoLshoot Cd translocation in *Arabidopsis thaliana*. New Phytol. 2009;181:71–8. 10.1111/j.1469-8137.2008.02638.x.

77. Xu G, Guo C, Shan H, Kong H. Divergence of duplicate genes in exon–intron structure. Proc Natl Acad Sci. 2012;109:1187–92. 10.1073/pnas.1109047109.

78. Xu E, Liu Y, Gu D, Zhan X, Li J, Zhou K, et al. Molecular Mechanisms of Plant Responses to Copper: From Deficiency to Excess. Int J Mol Sci. 2024;25(13):6993. doi:10.3390/ijms25136993.

79. Yin L, Wu R, An R, Feng Y, Qiu Y, Zhang M. Genome-wide identification, molecular evolution and expression analysis of the B-box gene family in mung bean (Vigna radiata L.). BMC Plant Biol. 2024;24:532. 10.1186/s12870-024-05236-9.

80. Yin L, Zhang M, Wu R, Chen X, Liu F, Xing B. Genome-wide analysis of OSCA gene family members in Vigna radiata and their involvement in the osmotic response. BMC Plant Biol. 2021;21:408. 10.1186/s12870-021-03184-2.

81. Zaun HC, Shrier A, Orlowski J. N-Myristoylation and Ca2+ Binding of Calcineurin B Homologous Protein CHP3 Are Required to Enhance Na+/H+ Exchanger NHE1 Half-life and Activity at the Plasma Membrane. J Biol Chem. 2012;287:36883–95. 10.1074/jbc.M112.394700.

82. Zhang C, Yang Q, Zhang X, Zhang X, Yu T, Wu Y, et al. Genome-Wide Identification of the HMA Gene Family and Expression Analysis under Cd Stress in Barley. Plants. 2021;10:1849. 10.3390/plants10091849.

83. Zlobin IE. Current understanding of plant zinc homeostasis regulation mechanisms. Plant Physiol Biochem. 2021;162:327–35. 10.1016/j.plaphy.2021.03.003.

84. Zorrig W, Abdelly C, Berthomieu P. The phylogenetic tree gathering the plant Zn/Cd/Pb/Co P1B-ATPases appears to be structured according to the botanical families. C R Biol. 2011;334:863–71. 10.1016/j.crvi.2011.09.004.

